# The first complete mitochondrial genome of tjaku□a (Great Desert Skink, *Liopholis kintorei*)

**DOI:** 10.1101/2024.07.17.603866

**Authors:** David Thuo, Jesse Wallace, J. Scott Keogh, Nicholas A. Macgregor, Margarita Goumas, Shaeleigh Swan, Tracey Guest, Erik D. Doerr, Rachel Paltridge, Jeremy Kenny, Samuel D. Merson, Leo Joseph

**Affiliations:** Australian National Wildlife Collection, CSIRO National Research Collections Australia, Canberra, ACT, Australia; Division of Ecology and Evolution, Research School of Biology, The Australian National University, Canberra, Australia; National Collections & Marine Infrastructure, CSIRO, Parkville, VIC, Australia; Parks Australia, Canberra, ACT, Australia; Ulu□u-Kata Tju□a National Park, Parks Australia, Northern Territory, Australia; Indigenous Desert Alliance, Alice Springs, Northern Territory, Australia; Mu□itjulu Tjaku□a Rangers, Mu□itjulu, Northern Territory, Australia; Durell Institute of Conservation and Ecology, School of Anthropology and Conservation, University of Kent, Canterbury, UK

## Abstract

The complete mitochondrial genome (mitogenome) of the tjaku□a, *Liopholis kintorei* was obtained using next-generation sequencing, making it the first recorded mitogenome of the genus *Liopholis* and the *Tiliquini*. The mitogenome is 16,844bp in length with a base composition of A (31.7%), T (24.4%), G (14.6%), and C (29.3%) and a G + C content of 43.9%. The genome contains 13 protein-coding genes, 22 transfer RNA genes, two ribosomal RNA genes (12S and 16S), and three non-coding fragments, consisting of the putative control region and two mitochondrially encoded heavy strand origin of replication region (OriH). The gene order is identical to that of typical skink mitogenomes. This genomic resource will provide valuable information for genetic studies of this genus and contribute to the growing collection of mitogenomes within the family Scincidae.

## Introduction

Scincidae is the most diverse Family within the Order Squamata, having approximately 1,760 described species (Uetz et al., 2023). Australia hosts a significant portion of this diversity in its over 400 species (ASH, 2023), the vast majority of which are endemic to the continent (ASH, 2023; Uetz et al., 2023). The genus *Liopholis*, recently separated from *Egernia* (Gardner et al., 2008), comprises 11 species distributed across Australia’s temperate and arid regions. Among them, *Liopholis kintorei* (Figure 1), locally known as tjaku a and culturally significant to the Aboriginal people of Central Australia (IDA, 2022), is one of the most threatened species. It is classified as ‘Vulnerable’ to (EPBC, 1999; IUCN, 2024) and was recently identified as one of Australia’s top 110 priority species requiring urgent conservation actions (DCCEEW, 2022)

**Figure 1.**
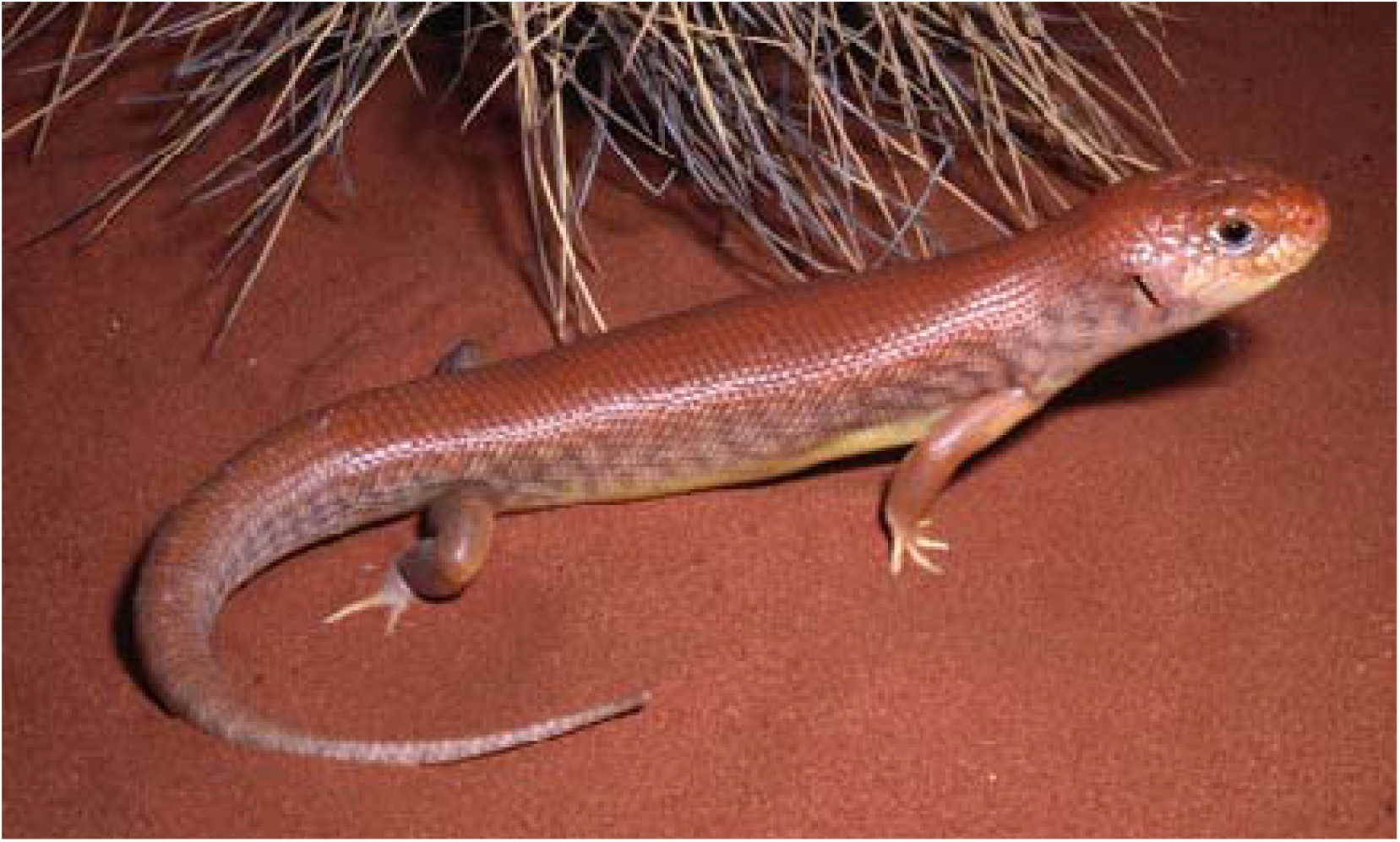
Photograph of tjaku□a, *Liopholis kintorei* taken by Steve Wilson. The picture was retrieved (June 15, 2024) from https://www.dcceew.gov.au/environment/biodiversity/threatened/action-plan/priority-reptiles/great-desert-skink

Despite its biological and cultural importance, scant molecular data from *Liopholis kintorei* have been published in the NCBI (Chapple & Keogh, 2004; Dennison et al., 2015). To address this gap and expand the genomic resources for this species, we present the first complete mitochondrial genome of tjaku□a. This mitochondrial genome of *Liopholis kintorei* will increase our understanding of the evolutionary history and phylogenetic relationships within the genus *Liopholis*, thereby informing the conservation strategies needed to safeguard tjaku□a.

### Mitochondrial genome assembly and annotation

A liver sample collected from a roadkilled specimen of tjaku□a (Australian National Wildlife Collection R13082) was used for this study. Total genomic DNA was extracted and sequenced using a next-generation sequencing platform. Prior to assembly, raw sequences were quality-trimmed with Trimmomatic (Bolger et al., 2014) and dereplicated using BBMap (Bushnell et al., 2017). High-quality reads were assembled using SPAdes (Bankevich et al., 2012) and mitochondrial contigs were identified using CONSULT-II (Şapci et al., 2024) with a RefSeq (O’Leary et al., 2016) mitochondrial genome database. Identified contigs were circularised using a custom Python script and annotated for tRNAs with MITOS (Bernt et al., 2013). Putative contigs were annotated for mitochondrial genes and reformatted to conform with HGNC gene nomenclature (Seal et al., 2020). Each contig was searched against the NCBI nt database (Sayers et al., 2020) using BLAST+ (Camacho et al., 2009) and only the top hit was used to ensure that the sequence did not arise from contamination. Final contigs were selected based on gene completeness, length, top BLAST hit, and successful circularisation. Homology with sequences from other conspecific samples was confirmed by aligning the selected contigs using MAFFT (Katoh & Standley, 2013). The OGDRAW (Greiner et al., 2019) was used to create a circular display of the tjaku□a, *Liopholis kintorei* mitogenome (Figure 2).

**Figure 2.**
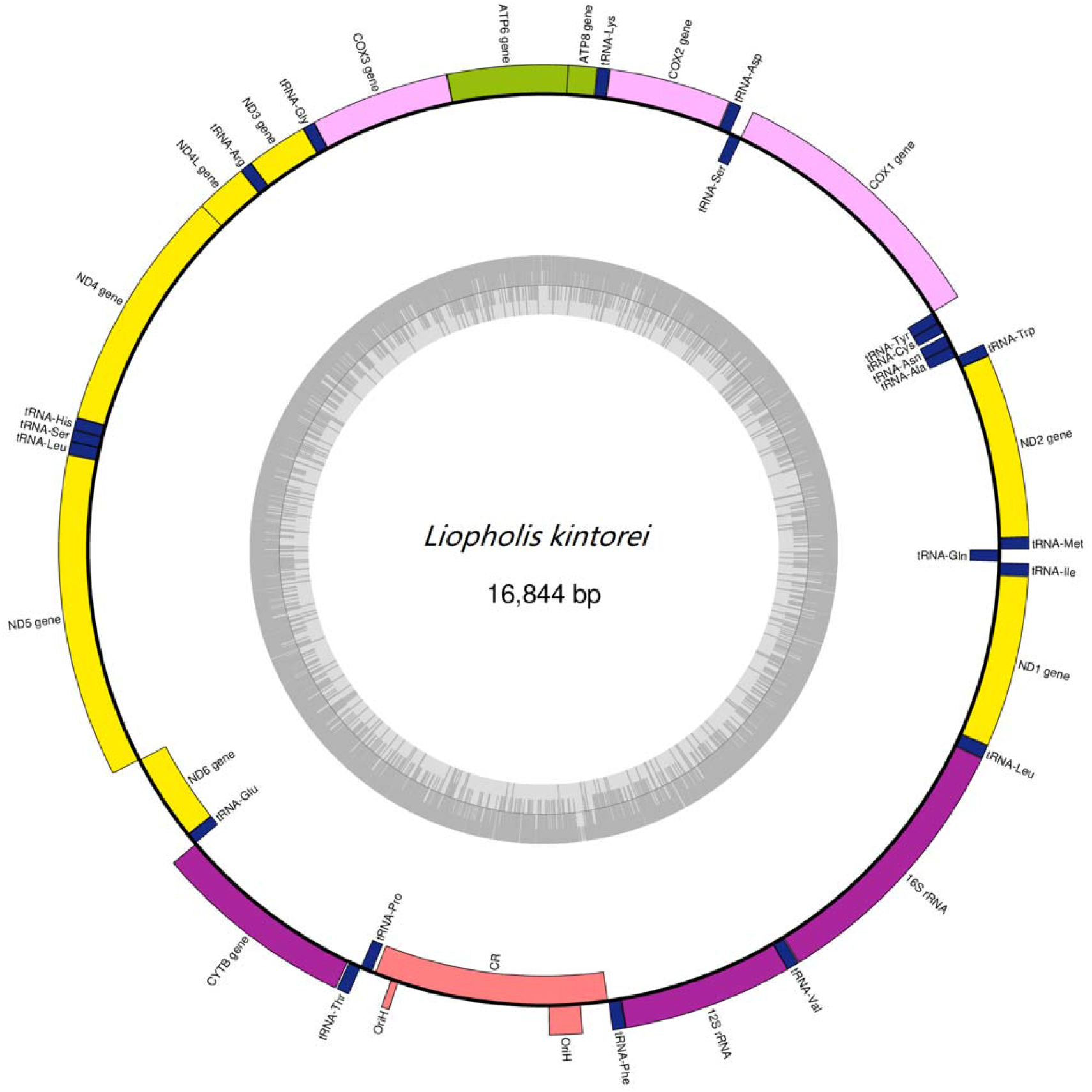
Circular visualisation maps of the complete mitochondrial genome of tjaku□a, *Liopholis kintorei*. The genes inside the map are encoded on the light strand.

## Results

The size of the tjaku□a mitochondrial genome is 16,844 bp in length (Figure 2; Table 1). It contains 22 tRNA genes, 13 protein-coding genes, 2 rRNA genes (12S rRNA and 16S rRNA), and three non-coding fragments (one control regional and two OriH). The synteny of all these genes is well conserved in comparison with the typical mitogenome of Scincidae (Chen et al., 2021, 2020; Park et al., 2016; Song et al., 2016; Zhong et al., 2021). The total base composition is A (31.7%), T (24.4%), G (14.6%), and C (29.3%) with a GC of 43.9%. The 12S rRNA (958 bp) and the 16S rRNA (1537 bp) are located between tRNA-Phe and tRNA-Leu but are themselves separated by tRNA-Val (Figure 2).

**Table 1.** Features of tjaku□a *Liopholis kintorei* mitochondrial genome.

| Feature | Sequence location | size | Intergenic region |
| --- | --- | --- | --- |
| tRNA-Met | 1-69 | 69 | 0 |
| ND2 | 70-1104 | 1035 | 0 |
| tRNA-Trp | 1104-1173 | 70 | 0 |
| tRNA-Ala | 1173-1241 | 69 | 0 |
| tRNA-Asn | 1242-1314 | 73 | 9 |
| tRNA-Cys | 1334-1398 | 65 | 0 |
| tRNA-Tyr | 1399-1467 | 69 | 1 |
| COX1 | 1469-3016 | 1548 | 3 |
| tRNA-Ser | 3012-3082 | 71 | 3 |
| tRNA-Asp | 3086-3154 | 69 | 2 |
| COX2 | 3157-3844 | 688 | 0 |
| tRNA-Lys | 3845-3911 | 67 | 1 |
| ATP8 | 3913-4077 | 165 | -8 |
| ATP6 | 4068-4751 | 684 | 0 |
| COX3 | 4751-5535 | 785 | 0 |
| tRNA-Gly | 5535-5602 | 68 | 0 |
| ND3 | 5603-5950 | 348 | 0 |
| tRNA-Arg | 5949-6017 | 69 | 0 |
| ND4L | 6018-6311 | 294 | 0 |
| ND4 | 6305-7682 | 1378 | -5 |
| tRNA-His | 7683-7751 | 69 | 0 |
| tRNA-Ser | 7752-7818 | 67 | 0 |
| tRNA-Leu | 7818-7890 | 73 | -1 |
| ND5 | 7892-9706 | 1815 | 1 |
| ND6 | 9707-10231 | 525 | 0 |
| tRNA-Glu | 10232-10299 | 68 | 0 |
| CYTB | 10302-11444 | 1143 | 2 |
| tRNA-Thr | 11452-11524 | 73 | 7 |
| tRNA-Pro | 11525-11595 | 71 | 0 |
| Control region | 11616-13012 | 1397 | 0 |
| tRNA-Phe | 13014-13088 | 75 | 2 |
| 12S rRNA | 13089-14046 | 958 | 0 |
| tRNA-Val | 14044-14113 | 70 | 1 |
| 16S rRNA | 14116-15652 | 1537 | 2 |
| tRNA-Leu | 15654-15728 | 75 | 1 |
| ND1 | 15730-16695 | 966 | 2 |
| tRNA-Ile | 16698-16769 | 72 | 2 |
| tRNA-Gln | 16775-16844 | 70 | 5 |

All protein-coding genes initiate with ATG as a start codon, except the COX1 gene which initiates with GTG. Nine protein-coding genes (ND1, ND2, ND3, ND4L, ND5, ATP6, ATP8, COX1 and CYTB) terminate with TAA, TAG or AGA stop codons. The remaining four genes (COX2, COX3, ND4 and ND6) have incomplete stop codons ending with either T or TA, which are presumably polyadenylated to a TAA stop codon (Ojala et al., 1981). ATG is the most common start codon (12 genes), and TAA is the most common termination codon (7 genes). The 22 tRNA genes are interspersed between rRNA and protein-coding genes, their lengths ranging from 65 bp to 75 bp (Table 1). Simulation analysis using tRNAscan-SE indicates that all tRNA genes in the tjaku□a mitochondrial DNA can potentially fold into typical cloverleaf structures, except for tRNA-Ser (Figure 3). The tRNA-Ser gene lacks the D-arm, and this has been observed in other members of the Scincidae (Chen et al., 2021). The major non-coding region in the tjaku□a mitochondrial DNA includes the Origin of Replication sequence and is 1397 bp long. The non-coding region is located between the tRNA-Pro and RNA-Phe genes. The conserved sequence within this sequence is similar to the conserved sequences found in other species of the family Scincidae.

**Figure 3.**
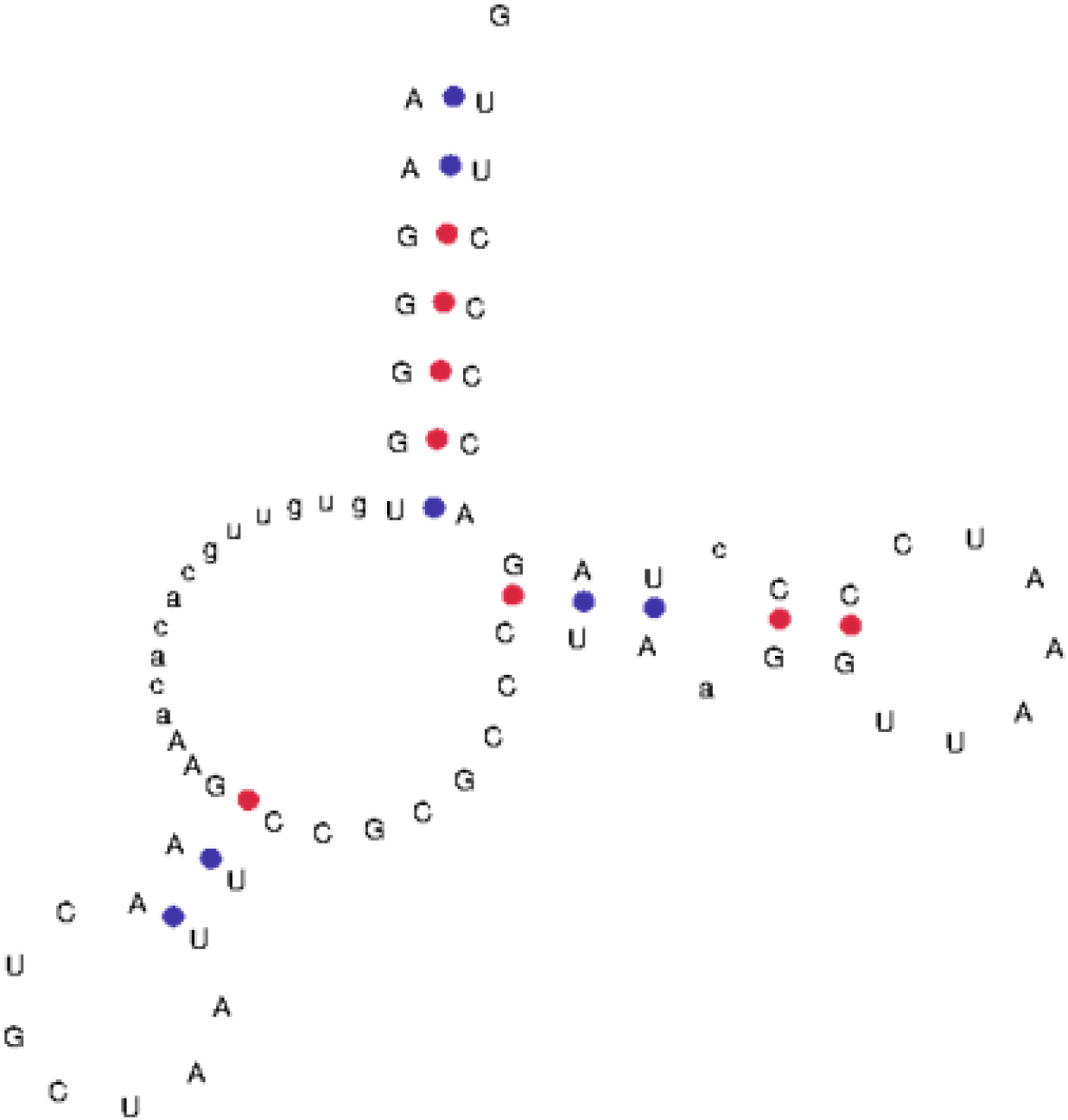
The predicted structure of tRNA-Ser lacks the D-Arm. Blue and red dots between A-U/G-C pairs indicate standard RNA base pairings.

## Conclusions

We report the first complete mitochondrial genome of tjaku□a (the great desert skink, *Liopholis kintorei)*. The genome is 16,844 bp in length with gene order and structure similar to other members of the family Scincidae. This mitochondrial genome is available in the GenBank database under the accession number PP957932.

## Acknowledgements

We thank Anna Kearns and Jenny Giles from the CSIRO’s National Biodiversity DNA Library (NBDL) for their assistance with DNA sequencing. We thank the Anangu traditional owners, Director of National Parks (Permit number: R1/22), Central Land Council (Authority number: P78658), Voyages Ayers Rock Resort and CSIRO Animal Ethics Committee (permit number: AEC 2022-01) for permitting us to conduct our study. This work was supported by the Commonwealth Scientific and Industrial Research Organisation (CSIRO), the Director of National Parks and National Parks Conservation Trust as part of the project titled ‘Molecules in the sand: eDNA, terrestrial environments, and the Biology of Tjaku□a, the Great Desert Skink *Liopholis kintorei’*.

